# Rolling out *plaque-2-sequence*: a single plaque sequencing approach enabling rapid, low-cost sequencing of phages directly from plaques

**DOI:** 10.1101/2025.11.01.684647

**Authors:** Slawomir Michniewski, Rebecca Glenny, Andrew Kinsella, Kane Porter, Kayne Wright, Sophie Harrison, Nzubechukwu Ugokwe, Amani Alrashidi, Srwa.J. Rashid, Chisomaga Eke, Jack C.D. Lee, Faizal Patel, Theo Josephs, Arezoo Pedramfar, Max Coleman, Hasanain F.Y Al-Dahash, Gibson A. Sabuholo, Navodya S. Roemer, Hannah Sampson, Gerald N. Misol, Branko Rihtman, David J. Scanlan, Elspeth Smith, Graham Stafford, Willem van Schaik, Richard J. Puxty, Edouard Galyov, Martha R. J. Clokie, Spyridon Megremis, Andrew Millard

## Abstract

Rapid, accurate, and scalable sequencing of bacteriophage genomes is critical to advance phage therapy, build phage biobanks and understand phage genomic diversity. Current methods are based on sequencing and assembling complete bacteriophage genomes using short- or long-read technologies. However, current protocols require large DNA input and are cost prohibitive which limits their application to phage collections that typically are large and have low-biomass.

In order to address this we have developed *plaque-2-sequence*, a robust and cost-effective workflow for high-throughput phage genome sequencing that will transform the speed and cost of attaining phage genomes. *Plaque-2-sequence* combines low-input transposase-based library preparation, amplification, nanopore sequencing and optimised assembly steps tailored to phage genomes. We applied the method to phages isolated on seven genetically diverse bacterial hosts; *Escherichia*, *Pseudomonas, Synechococcus, Enterococcus, Klebsiella , Serratia* and *Enterobacter.* High quality genome assemblies were validated using CheckV and benchmarking against previously sequenced phage isolates. Compared to standard Illumina sequencing, plaque-2-sequence offers ∼10-fold savings in sequencing price for individual labs. Furthermore, it substantially decreases the time required to produce a phage genome, once a plaque is obtained.

Offering the ability to routinely obtain hundreds of phage genome sequences a week, with minimal hands-on time. Plaque-2-sequence enables systematic genomic characterisation of phage isolates, facilitating taxonomic classification, for the development of large scale phage biobanks.

**Impact Statement:** Here we have optimised a method for high-throughput sequencing of bacteriophage genomes from single plaques (plaque-2-sequence). We present a robust, high-throughput and cost-effective workflow. Plaque-2-sequence combines low-input transposase-based library preparation, amplification, nanopore sequencing and optimised assembly steps tailored to phage genomes. We demonstrate the scalability of this approach by sequencing over 100 phages from multiple bacterial hosts. This marks a step-change for the field, allowing phage genome sequencing to keep pace with phage isolation rates, and transforming how rapidly we can explore and understand phage genomic diversity.

## Introduction

Bacteriophages, also known as phages, are viruses which selectively target and kill bacteria. All phages consist of a protein or proteolipid capsid enveloping the viral genetic material [1]. Although phages exhibit strong morphological similarities and conservation of structural proteins leading to a small number of known morphotypes [2], their genomic diversity is vast [3]. To date the vast majority of isolated phages belong to the tailed group of dsDNA phages of the class Caudoviricetes [4]. Non-tailed dsDNA phages such as *Corticoviridae [5]*, ssDNA phages such as filamentous *Inoviridae* [6] or icosahedral *Microviridae [7]* and RNA phages of the class *Leviviricetes* are also known [8], but are under-represented in genomic databases.

Unlike bacteria where the 16S rRNA gene is commonly used as a conserved marker gene for taxonomic assignment, bacteriophages have no universal conserved genes that can be utilised for rapid genomic assessment. Several sets of signature genes such as *polA , polB, g20* and *g23* have been used where prior knowledge of the expected phage type exists [9]. However, the only way to determine if a newly isolated phage is different from previously described phages, is to sequence its genome. Phage genome sequencing is commonplace, with >30,000 genomes from cultured phage isolates in public databases [3]. To date, the vast majority of these have been sequenced with Illumina sequencing methods [3]. Long read sequencing methods such as Oxford Nanopore Technology (ONT) and PacBio, have also been used to sequence phage genomes and metagenomes [10–12].

With general renewed interest in phages, and specific interest in phage therapy, there is an increasing need for high-throughput phage genome sequencing. In the context of phage therapy, particularly during the discovery phase, sequencing large numbers of phages is essential to capture their full genetic diversity. This ensures that the most effective and therapeutically promising phages are rapidly identified, prioritised, and developed for clinical application. [13, 14]. The greater the initial diversity of the set of phages that are characterised, the greater the chances of identifying phages that have the desired characteristics. Furthermore, excluding phages that contain genes encoding for virulence factors, antimicrobial resistance or lysogeny associated proteins, can only be done by genome sequencing [15, 16]. The long turnaround times of commercial sequencing services combined with the need to generate high-titre phage stocks to obtain sufficient DNA, means that often preliminary phenotypic characterisation is carried out before genome data are available. This approach is inefficient, as phages may later be excluded due to redundancy or the presence of undesirable genes once sequencing is complete. There is therefore a pressing need for rapid, scalable, and cost-effective sequencing methods that enable early genomic screening of large phage collections, ensuring that only genetically distinct and therapeutically appropriate candidates progress to detailed characterisation.

Obtaining sufficient quantities of high-quality phage DNA remains one of the key bottlenecks in genome sequencing. Commercial sequencing suppliers typically require hundreds of nanograms of DNA [17], which means high-titre phage stocks and at least millilitre-scale lysates [18–20]. Although this is fine for a few isolates, it is not sustainable when hundreds of phages need to be processed in parallel. Illumina-based tagmentation protocols can, in principle, use a nanogram of DNA, but access to in-house sequencing facilities where such low limits can be flexibly applied is not readily available to most researchers. The Oxford Nanopore’s MinION platform offers a portable and cost-effective alternative, but similar DNA input requirements also limit the technology in its current form. Both platforms also share inherent limitations in that tagmentation-based library preparation excludes ssDNA phages, and many phage genomes contain base modifications that inhibit enzymatic library preparation [21–23]. Together, these factors slow our ability to systematically sequence and compare large, diverse phage collections.

Overcoming these barriers is critical if genome sequencing is to become the *starting point* of phage characterisation rather than its endpoint. A workflow that enables rapid, low-cost sequencing directly from small-scale lysates would allow early exclusion of redundant or undesirable isolates, incorporation of previously intractable phage types, and a truly diversity-led discovery process. In the context of phage therapy, this shift is transformative allowing genetic insight to guide isolation and formulation from the outset, rather than retrospectively. Our goal, therefore, was to develop a sequencing approach that is scalable, accessible, and cost-efficient, with minimal hands-on time and capital investment which will transform phage genomics from being a specialist bottleneck into a routine, enabling step change in therapeutic phage development.

## Materials and Methods

Bacterial and phage isolates

Bacterial strains and phages used in this study are detailed in Table 1

**Table 1.**
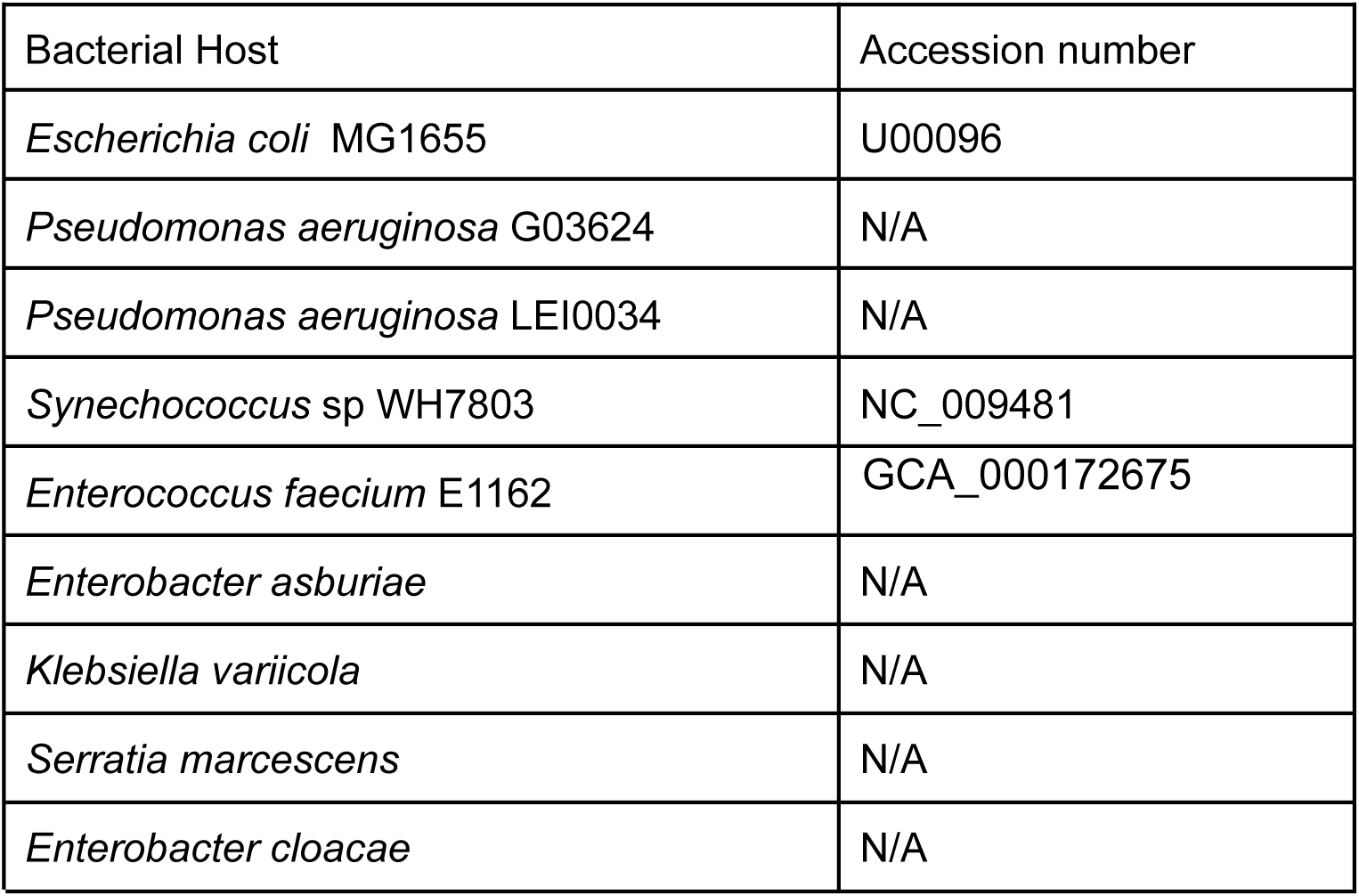
Bacterial hosts used in this study.

### Phage Isolation against *Pseudomonas aeruginosa and E. coli MG1655*

Plaque assays were performed using a standard protocol [17] with *Pseudomonas aeruginosa* as the host, using water from a pig trough. The resulting plaques were picked using a 1 mL pipette tip into separate 1.5 ml Eppendorf tubes containing 100 μL SM buffer (200 mM NaCl_2_, 10 mM MgSO_4_ , 50 mM Tris-HCl, pH 7.5), which was then vortexed and stored at 4 ℃ for further phage purification and/or propagation. A subsequent spot test using 10 μL of the previous sample was performed to confirm whether the plaques contained phages by the formation of a zone of clearing. Finally, a soft agar plug was collected from the middle of the resultant clearing zone using a 1 mL pipette filter-tip, transferred to a fresh 1.5 mL Eppendorf tube containing 100 μL SM buffer, vortexed and left for 1 hour for phages to disperse from the agar. Subsequently, the sample was centrifuged at 13 000 g for 5 minutes, and 10 μL of the supernatant transferred to a well in a 96 well plate. Phages against *Escherichia coli* MG1655 were collected in a similar manner, using water from Barston Sewage Works as the sample source.

### DNase I treatment prior to whole genome amplification

Each well within a 96 well plate was supplemented with 0.01 U of DNase I (Thermo Scientific) and incubated at 37 °C for 60 minutes followed by 95°C for 10 minutes, to ensure complete inactivation of the DNase I and simultaneous release of encapsidated phage DNA. Phage DNA samples were stored at -20 °C until further use.

### Whole genome amplification and debranching of the products

Whole genome amplification of phage DNA samples was performed in a separate 96 well plate using the EquiPhi29™ DNA Amplification Kit (Thermo Scientific) and following the manufacturer’s guidelines for standard 20 μL reactions, with reaction volumes reduced proportionally to half, one third or a quarter of the standard reaction. Debranching of amplified DNA was performed by supplementing each well in a 96 well plate with 10 U of S1 Nuclease (Thermo Scientific) containing buffer (1x final concentration) and incubated at 25 °C for 30 minutes followed by 95 °C for 10 minutes, to ensure complete inactivation of the S1 Nuclease. DNA concentration was quantified using a Qubit Broad Range kit (Thermo Fischer). Amplified phage DNA samples were stored at -20 °C until further use.

### Sequencing and genome assembly

Oxford Nanopore Technology (ONT) sequencing was carried out using the Rapid sequencing DNA V14 - barcoding kit (SQK-RBK114.24 or SQK-RBK114.96) on a MinION Mk1c/Mk1B or GridION, with R10.4.1 flow cells. Basecalling was carried out using the High-accuracy v4.3.0 - 400 bps model. Reads were trimmed for barcodes and adapters after sequencing, either within MinKnow or using dorado [18]. Long read assemblies were carried out with flye v2.8.1-b1676 using the following settings: “–interations 2, –nano_corr”. Followed by two rounds of polishing with medaka using the relevant model [18]. Genome assembly was carried out on the ALICE High Performance Computing facility at the University of Leicester, with 48 cores allocated for assembly.

### Genomic analysis

#### In silico dataset

Long reads were generated against a set of 196 phage genomes (Table S1), which were representative of all families of DNA viruses known to infect bacteria in the ICTV VMR_v40.1 . Long reads were generated using PBSIM3 [19] with the following settings “--strategy”, “wgs”, “--method”, “--errhmm”, model_path, “--depth”, “70”, “--length-mean”, “2500”, “--length-max”, “15000”, “--accuracy-mean”, “0.99”, “--accuracy-min”, “0.80”, “--length-sd”, “5000”. Short reads were then created from these long reads using a biopython script. Prior to assembly reads were normalised with <u>bbnorm.sh</u> target=150 [20]. Genomes were assembled with SPAdes v3.13.1 with “-s”, for single read libraries [21]. Assembled genomes were compared against the reference genome with dnadiff [22] to identify single nucleotide polymorphisms (SNPs) and insertions/deletions (INDELs). Completeness of genomes was determined by the formula:

> 𝐴𝑠𝑠𝑒𝑚𝑏𝑙𝑦 𝐿𝑒𝑛𝑔𝑡ℎ / 𝑅𝑒𝑓𝑒𝑟𝑒𝑛𝑐𝑒 𝑆𝑒𝑞𝑢𝑒𝑛𝑐𝑒 𝐿𝑒𝑛𝑔𝑡ℎ 𝑥 100

### Assembly Pipeline

Reads were basecalled using Dorado v 0.9.0 with “ basecaller hac --kit-name ” and fastq reads as output. Any chimeric reads were identified and split with SACRA v2.0 [23]. The resulting split reads were initially assembled with Flye v2.9.5 using “*nano_corrected*”. Assembled contigs were polished twice with Medaka v2.0.0 using “*medaka_consensus*”, using the relevant model. Phage genomes within each contig file were identified with CheckV v1.0.3 “end_to_end” to rapidly identify complete phage genomes [24]. All “Complete”, “High-Quality” and “Medium” genomes were extracted and taxonomy was assigned to complete and high-quality genomes using *taxMyPhage* [25].

### Short read assembly

Long reads were converted to short reads of length 300 bp with a python script. Assembly was then carried out as previously described [26], with minor modifications. Briefly, reads were normalised with bbnorm “--target=150”, prior to assembly with SPAdes v “-s , -- only-assembler”, allowing assembly with on single ends reads. All “Complete”, “High-Quality” and “Medium” genomes were extracted and taxonomy was assigned to complete and high-quality genomes using *axMyPhage*t [25]. Scripts used for genome assembly are available on github (https://github.com/amillard/plaque2seq).

### Quality control of sequencing data

The proportion of reads assumed to be host derived, was determined by mapping reads against the host genome with “minimap2 -ax map-ont -t {threads} {ref_fa} {input_fq} | samtools view -Sb - | samtools sort -o {output_bam}” . The number of mapped reads was determined with pysam [27] and calculated as a percentage of total reads. *Pseudomonas* prophages were predicted with geNomad “end-to-end” [28] and these regions manually excised from the host genome to create prophage free reference, which was then used to calculate the percentage of mapped host reads.

### Genome annotation,taxonomic classification and comparative genomics

Genomes were annotated with Pharokka v1.7.5 [16] using PHROGS HMMs [29]. All genomes were submitted to the European Nucleotide Archive (ENA), see supplementary tables for the accession numbers (Table S2). Bacteriophages were classified into existing genera and species utilising Iv0.3.3, using default settings [25]. Bacteriophages were compared to each other using the *taxMyPhage* “similarity” option, with genomes identified as 100% ANI, classified as identical.

### “Warwick” and “Sheffield” , phage isolation, genome sequencing and assembly

To validate the approach, the protocol was carried out by researchers at the University of Warwick and Sheffield

### Phage isolation against *Synechococcus* sp. WH7803

Cyanophages were isolated from seawater samples against *Synechococcus sp*. WH7803 according to standard protocols [30]. Initially, 200 µL of seawater sample from the Atlantic Meridional Transect 24 (AMT24) research cruise was inoculated into 1.8 mL *Synechococcus* sp. WH7803 culture and incubated for two weeks at 22°C at a continuous light intensity of 10 μmol photons m^−2^ s^−1^. The infected cultures were then centrifuged at 4,000 x g for 10 minutes and filtered through 0.2 µM pore size filters to obtain the phage fraction. Subsequently, 20 µL aliquots of the phage fraction were spotted onto *Synechococcus* sp. WH7803 lawns to screen for phage-producing samples. Samples which produced a zone of clearing were resuspended in 1 mL ASW medium [30]. To obtain clonal phages using plaque assays, 10 µL of the resuspension was added to 100 µL 10x concentrated *Synechococcus* sp. WH7803 and incubated for one hour at 22°C with illumination. The infection was then mixed with 3 mL cooled molten ASW medium agar (0.2% w/v), and incubated as above. After incubation, single plaques were picked and resuspended in 1 mL ASW medium. This process was repeated twice more to obtain clonal cyanophages. Clonal cyanophages were subsequently used for sequencing as described above, with the omission of the S1 nuclease step. Phage genomes were sequenced on a MinION Mk1C. Reads were base called using Dorado v 0.9.0 with “ basecaller hac --kit-name ” , with fastq reads as output. Genomes were assembled from short reads as described above, and resulting genomes were annotated with Pharokka using the PHROGs database, as previously described [26, 29, 31]

### Phage isolation against *Entrobacter, Enterococccus ,Klebsiella* and *Serratia*

Phages were isolated from wastewater samples collected from Blackburn Meadows, Sheffield or animal faeces donated by Yorkshire Wildlife Park, Doncaster. 150 µl of filtered (0.45 µm pore size) environmental samples were mixed with 0.5 ml of bacterial culture (OD 0.5-1) and incubated for 10 minutes at room temperature. After incubation, 5 ml of BHI soft agar (0.4% w/v) (Avantor, USA) was added to the mixture, inverted to mix, and then immediately poured on top of a solid BHI agar plate and incubated overnight at 37°C. Plaques were then picked and resuspended into 50 µl of PBS and stored at 4°C until re-infection. To ensure the purity of the possible phage, three successive rounds of plaque picking and plaque assays were carried out. For testing *plaque-2-sequence,* a single plaque was picked and resuspended in 100 µl of PBS, vortexed and left for 1 hr to resuspend. The plaque resuspensions were then used for sequencing as described above, with the omission of the S1 nuclease step.

Reads were basecalled with Dorado v 0.9.0 with “ basecaller sup --kit-name ” , with fastq reads as output. The phages Klebsiella phage Kv_Kong and Enterococcus phage Efm_George were sequenced by both *plaque-2-seq* and standard extraction of DNA from high titre lysate (un-amplified). For reads generated from un-amplified DNA , assemblies were carried on the Galaxy web platform (https://usegalaxy.eu) . Fastq reads were assembled using Raven v1.8.3 with default parameters [32] , assembled contigs were polished with the Medaka Consensus Pipeline v1.7.2 using the model ‘r1041_e82_400bps_sup_g615’ (https://github.com/nanoporetech/medaka). The resulting contig(s) were analysed with CheckV v1.0.3 to identify complete phage genomes [24]. Genomes were annotated with Pharokka v1.3.2 [16]. Phage SM58.13_P2 was also assembled using the approach described above. All genomes were assembled by the previously described approach of creating short short reads from long reads, with the exception of Serratia phage SM58.13_P2 which did not produce an assembled genome from short reads. Host contamination was calculated by mapping of reads to the complete phage genome with Minimap2 v2.28 [33]. Host reads were calculated as those that did not map to the phage genome.

## Results

To remove the labour intensive step of multiple rounds of plaque purification and production of high-titre stocks for DNA extraction, we tested the use of whole genome amplification using multiple displacement amplification (MDA) of phage DNA to provide sufficient DNA for sequencing from a single plaque. As the aim was to develop a process which was as cost-effective as possible, we tested 0.5x, 0.3x and 0.25x volumes relative to the 20 µL recommended volume by the manufacturer of EquiPhi29 MDA kits, as well as testing 1, 2 and 4 hour incubation times with a single plaque as input material. There was no significant difference in yield of DNA obtained, under the nine conditions tested with all producing sufficient DNA for sequencing (Figure S1). Thus, 0.25 volumes of the recommended amount was used in further experiments (Figure S1).

Having established DNA could be amplified from a single plaque, we next determined if different treatments were required to successfully assemble phage genomes from the obtained DNA. As the process was based solely on touching a plaque with a sterile tip, there is a high probability of carry-over of host DNA. Furthermore, the process of MDA amplification is known to produce chimeric branching into DNA [23]. Thus, we designed an experiment whereby we could assess if these factors might impact our results. In short, 48 single plaques were obtained, using a single water sample from a pig trough, against a clinical isolate of *Pseudomonas aeruginosa*. All plaques were resuspended in 100 μl of SM buffer and treated using four different protocols, prior to sequencing (Table 2).

**Table 2.** Treatments applied to 48 plaques obtained against *Pseudomonas aeruginosa*.

|  | DNase I<br>Treatment | Amplification | S1 Nuclease | Gb of Data |
| --- | --- | --- | --- | --- |
| Treatment 1 | N | Y | N | 9.65 Gb |
| Treatment 2 | Y | Y | N | 2.84 Gb |
| Treatment 3 | N | Y | Y | 7.53 Gb |
| Treatment 4 | Y | Y | Y | 9.14 Gb |

The chosen method of sequencing was Oxford Nanopore Technology, due to the minimal capital costs associated with the equipment. Of the expected 192 samples, DNA was obtained from 174 samples, with failed samples due to edge effects associated with incomplete sealing of 96-well PCR plates. For each treatment group, one sequencing library was produced, with up to 48 barcodes per flow cell. Each library was sequenced on a different R10.4.1 flow cell.

## Sequencing results

The total amount of sequencing data produced per treatment (174 plaques) varied from 2.84 to 9.65 Gb of data (Table 1). The largest yield of sequencing data was obtained from treatment group 1 (9.65 Gb), closely followed by groups 4 ( 9.14 Gb) and 3 (7.53 Gb) , with data output from treatment 2 (2.84 Gb) being the lowest. As MDA is known to introduce chimeric reads [23], we calculated the percentage of reads that contained putative chimeras for each treatment. In treatments 1 and 2, during which no S1 nuclease was applied, the median percentage of chimeric reads was 60.2 and 62.7 (Figure 1). There was a significant decrease (Mann-Whitney U test , with Benjamini–Hochberg correction p < 0.05 ) in the percentage of chimeric reads in treatments 3 and 4 with 48.5 and 49.9 respectively. Thus, the inclusion of S1 nuclease in our protocol reduced the number of chimeric reads, but did not eliminate all chimeric reads.

**Figure 1.**
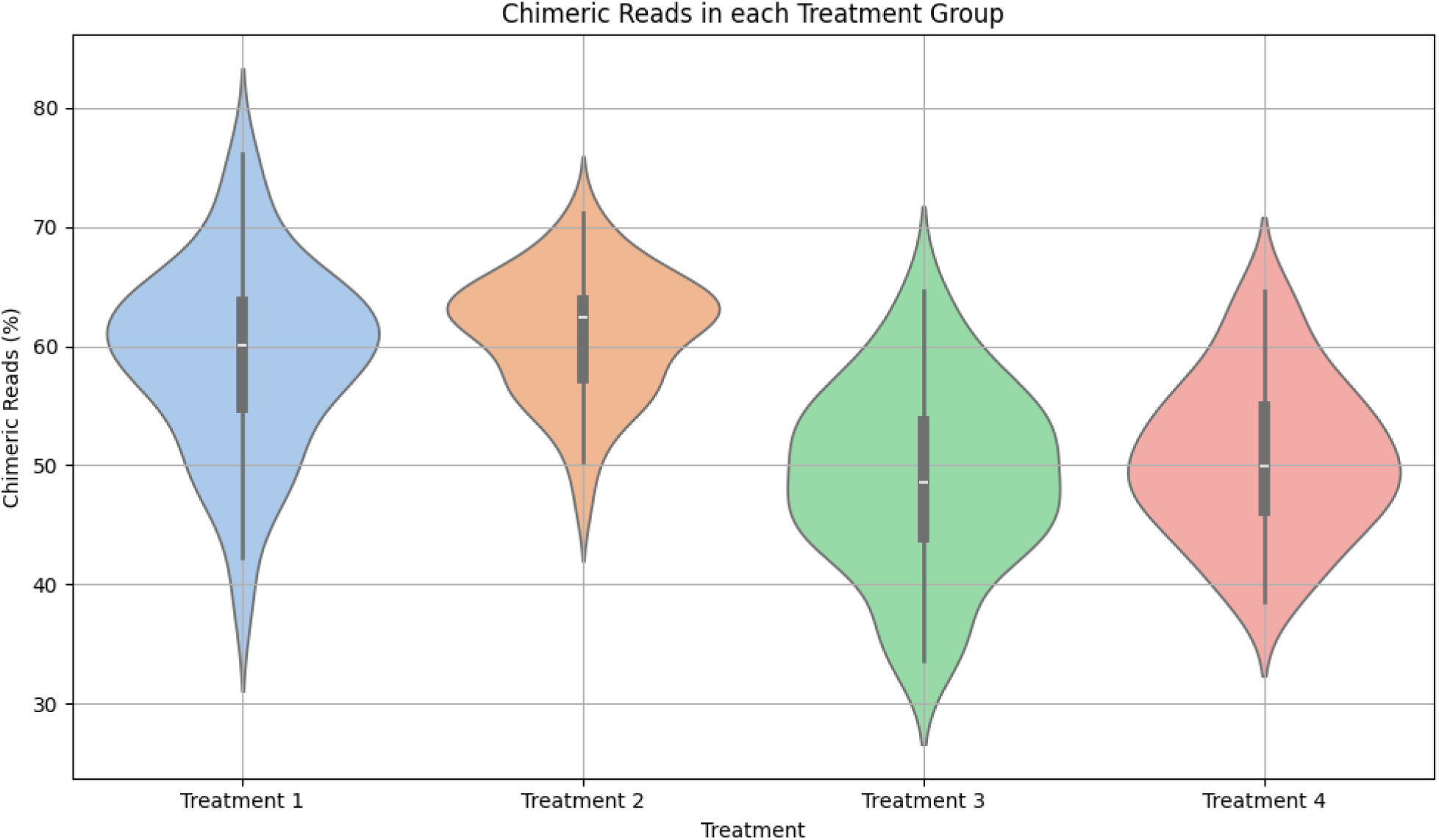
Detection of chimeric reads in four test conditions. The percentage of chimeric reads was determined from the output of SACRA [23] . Treatment 1:MDA only Nuclease; Treatment 2: MDA and DNAse; Treatment 3: MDA and S1 Nuclease; Treatment 4: MDA, S1 Nuclease and DNase I.

To determine the effect of DNase I treatment, reads were mapped against the host *Pseudomonas aeruginosa* reference genome sequence to calculate the percentage of contaminating host reads. However, >80 samples were found where >90% of reads mapped to the host genome, suggesting no phages were present, high levels of induced prophages or a combination of both. To overcome this, prophage regions were predicted with geNomad in the host genome and these regions excluded from inclusion in the calculation of reads mapped to the host genome. However, a high percentage of reads were still found across all samples. With no significant differences in the percentage of reads mapped to the host genomes across the four treatment groups (Figure S2). The optimal conditions for DNA amplification and sequencing was utilising 0.25x volumes of MDA reagents with a S1 nuclease step. As DNAse I wasn’t detrimental and has minimal cost, we included this in further work.

## Bioinformatics optimisation

We next sought to automate the assembly of phage genomes. We utilised Flye for the assembly of long reads, polishing with medaka, and checkV for the automated identification of phage genomes within each sample (Figure 2). We compared this process with an additional step of utilising SACRA for the identification and splitting of chimeric reads [23], prior to assembly. Although 48 plaques were selected, due to the aforementioned failed DNA amplification or failed sequencing, only 39/48 samples were present in all treatment groups. Only samples present in all four treatment groups were included in the comparative genome analysis. The number of complete, high-quality and medium-quality genomes was assessed with checkV.

**Figure 2.**
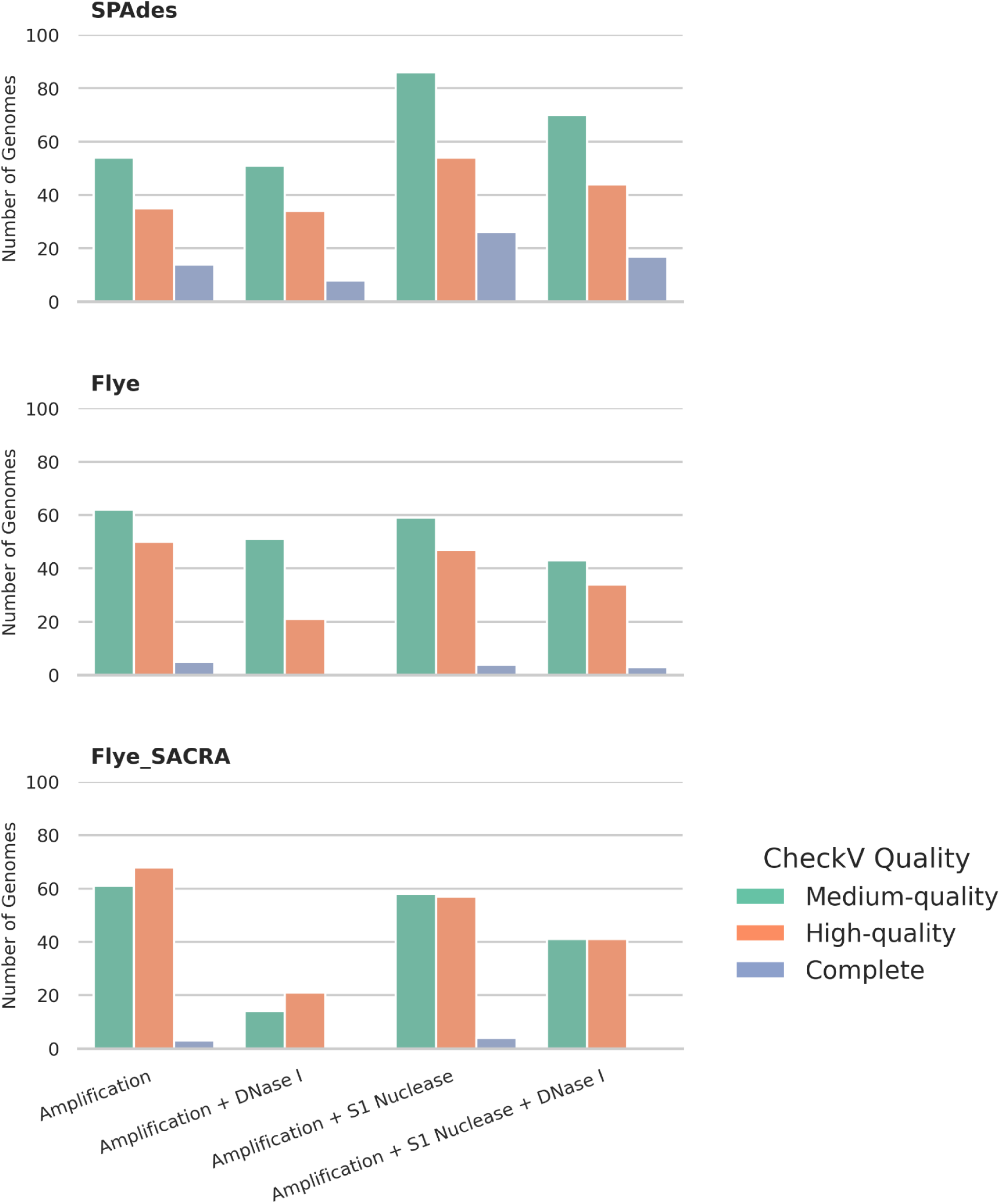
Comparison of assembly approaches. Three different assembly approaches were compared: SPades - assembly of short reads; Flye - assembly of long reads; Flye+SACRA - assembly of long reads with detection and splitting of chimeric reads with SACRA[23]. The number of phage genomes in each treatment group was determined using CheckV[24]. Treatment 1: MDA, S1 nuclease only; Treatment 2: MDA and DNase I only; Treatment 3: MDA and S1 nuclease only; Treatment 4: MDA, S1 nuclease and DNase I.

Across all conditions, the use of Flye+SACRA generally increased the number of contigs that could be identified as phage within the assemblies (Figure 2). A comparison of the ‘Flye alone’ and Flye+SACRA assembly approaches, on samples where no DNase I or S1 nuclease step was applied (Treatment 1), showed that a greater number of predicted complete and high-quality genomes were observed utilising SACRA to split chimeric reads (Figure 2). A similar pattern was observed even when both S1 nuclease and DNase I were utilised (Treatment 4), with the combined number of high quality and complete genomes increasing from 37 when using Flye alone to 41, when utilising Flye+SACRA (Figure 2). While SACRA was useful in the identification of chimeric reads, and resultant assemblies, it had an additional cost of significantly increased computational time. The assembly of 48 genomes when utilising SACRA was ∼36 hours, compared to < 12 hours without.

To reduce the time for assembly, we hypothesised that turning long reads into short reads and using a short read assembler would decrease the assembly time. We first tested if by creating short 300 bp reads from long reads, and then assembling with the short read assembler SPades would allow recovery of complete phage genomes. To test the applicability of this approach we first created a simulated set of long reads, from a representative set of bacteriophage genomes (Table S1), that were then made into short reads. The percentage completeness (compared to the original reference) for this in silico dataset ranged from 83.39 to 100.1% , with a median of 99.79% (Figure S3) thus demonstrating the approach works for simulated reads.

When applied to the data generated in this study, the largest number of complete and high-quality genomes in all conditions was observed when utilising SPAdes (Figure 2). Complete and high-quality predictions have been combined for comparison, as CheckV utilises terminal repeats to classify complete genomes from high-quality. However, SPAdes introduces repeated kmers at the ends of circularly permuted contigs, which are classified as complete, whereas Flye does not introduce such kmers to identify circular contigs. Any circularly permuted genomes will be called complete if assembled by SPAdes, but not Flye, even if both are complete.

The automated assembly and phage identification approach identified many more than the 39 genomes which would be expected if each plaque represented a single phage (Figure 2), with 80 genomes in treatment group 3 using SPAdes assembly. The reason for this is the assembly (and partial assembly) of prophages from the host organism, which were assembled at the same time as any isolated phages. Prophages were identified by comparison against the host genome. Of the original 48 plaques across all four treatments, it was possible to determine 19 solely contained prophages, suggesting they were caused by spontaneous induction of host prophages. Four plaques contained two phages, other than prophages, and 21 plaques were caused by other phages (although still contained prophage DNA). Thus, this approach allowed the rapid identification of 29 new phage isolates.

### Scaling up to 96 barcodes

Having established short reads, derived from long reads, can be utilised for single plaque sequencing, we further tested the approach with *E. coli* MG1655 as a host, increasing the number of plaques tested to 96 from a single water sample. We utilised both an S1 nuclease step as this reduced chimeric reads, and DNAse I treatment in library preparations. Of these 96 samples, 95 produced sufficient DNA for further sequencing. The approach of MDA was remarkably consistent with a median yield of 443 ng/µL (+/- 146 ng). As *E. coli* MG1655 has only cryptic prophages [34], the levels of host DNA per library were readily quantifiable by read mapping to the host chromosome. The median host DNA contamination was 2.75% (+/- 19%) per sample. Sequencing yield per sample varied across samples, with no correlation of input amount and output yield (Figure 3). Despite this variability, it was sufficient to provide the depth of sequencing required for a single phage genome. Assuming a mean phage genome size of 75 kb, the median predicted depth of sequencing would be 302x per phage genome, which far exceeds the ∼30x coverage needed for genome assembly [35] .

**Figure 3.**
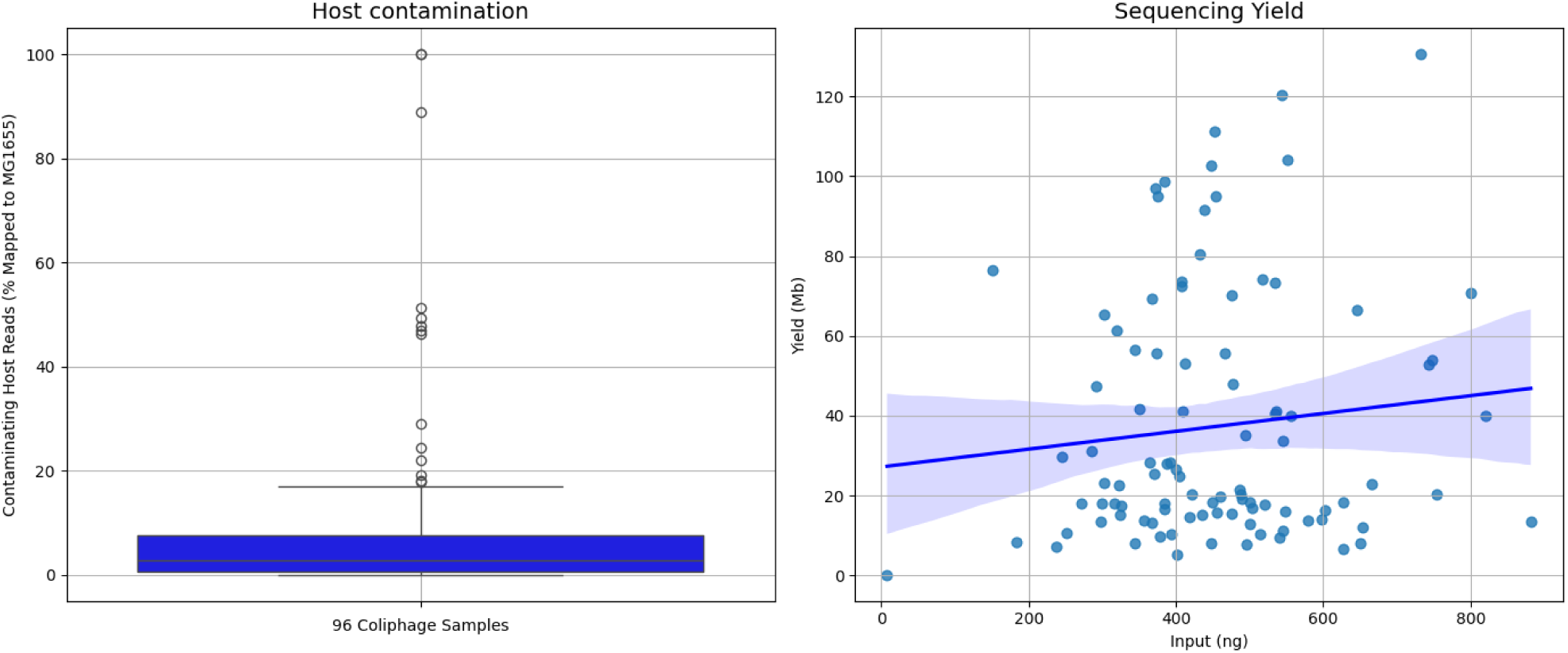
Output of sequencing data for 96 coliphage samples. A) Boxplot of host contamination for 96 coliphage samples. Host contamination was determined by mapping of reads to *E. coli* MG1655. B) Sequencing yield (Mb) plotted against library input (ng) for 96 coliphage samples.

The resulting assembly of 87 samples into 91 genomes confirms sufficient coverage was obtained. For samples which did not result in the production of a phage genome (n = 9), one of these was a result of insufficient DNA amplification and two from amplification of solely bacterial DNA. The failure of the other six samples was not clearly identifiable. Four samples were found to have two phages present. The rapid sequencing of plaques from a single source material, also rapidly identified the diversity of phages present within a single sample. Of the 91 coliphage genomes assembled, 42 of the 91 were unique when compared against each other (Figure S4). From this single source of wastewater used for isolation, phages classified into two families (*Drexlerviridae, Straboviridae*) and seven genera were isolated (*Dhillonvirus, Felixounavirus, Hanrivervirus, Krishvirus, Tequatrovirus, Vequnitavirus and Warwickvirus* ) (Table S4).

### Genome fidelity

In order to test the fidelity of the process, we resequenced the *Salmonella* phage RA112 using nanopore reads, and compared this to the original Illumina assembly, resulting in an identical genome sequence. Furthermore, we sequenced three plaques from the same coliphage (CCE_0049) and obtained identical sequences (Table S2). Repeating a similar process using Pseudomonas phage GAS_0134 produced identical results, and the ssDNA *Pseudomonas* phage GAS_0135 also produced identical genomes, again confirming the reproducibility and accuracy of the approach.

### Independent validation

Having established the protocol at the Becky Mayer Centre for Phage Research at the University of Leicester, the approach was independently carried out at the University of Warwick and University of Sheffield. Using the approach a further 24 plaques obtained on Synechococcus sp. WH7803 were sequenced at the University of Warwick. A total of 21 genomes were obtained ranging in length from ∼171 to ∼226 kb. The failure of three genomes likely resulted from insufficient data, with three samples yielding < 5 MB of sequence data. Why a further two genomes did not assemble was unclear. The mean level of contaminating host DNA reads was estimated to be 0.825% +/- 2.6%, by mapping to the host genome.

A further seven phages were isolated and sequenced at the University of Sheffield isolated on *Enterococcus faecium, Enterobacter asburiae, Klebsiella variicola, Serratia marcescens* and *Enterobacter cloacae*, confirming the approach can be used on different bacterial species to sequence phage genomes that range in size from ∼150 kb to ∼275 kb. Phages Kv_Kong and Efm_George were sequenced using both amplified and un-amplified DNA, comparison of the resulting genomes using DNAdif [36] confirmed no differences between the two approaches. Thus, demonstrating the approach of plaque-2-sequencing works on plaques from a range of bacterial species and produces genomes of the same quality as other approaches. The level of host contamination was 7.19 % +/-8.42% across the samples sequenced.

## Discussion

We have developed an approach to rapidly sequence bacteriophage genomes that can be scaled to sequence thousands of bacteriophage genomes, at greatly reduced cost. By optimising each individual step we have developed a process which allows phage genomes to be sequenced for ∼£10 per genome, when applied to batches of 96 plaques. The process can also be applied to a smaller number of genomes in batches of 24 or 48 at an increased cost of ∼ £15, or individual phages with increasing cost. The approach does not require substantial capital costs, with the entry level MinION Mk1D available for < £3,000 (May 2025). We have substantially reduced the time taken to obtain complete phage genomes, streamlining the process such that 96 plaques can be “picked” on Monday with complete phage genomes obtained by Friday/Saturday by a single user.

The cost savings generated by our approach could be further improved by washing the flow cell between library runs. The theoretical output of a minION flow cell is 50 Gb of data. However, actual obtained output per flow cell is dependent on a number of factors [37]. Within this study we achieved ∼10 Gb of data from a flow cell on multiple occasions. Only 30x coverage is required for the assembly of phage genomes [35]. Assuming a genome size of 75 kb , with a 60x coverage requires 0.864 Gb of data for 192 phage genomes (with no host contamination). Thus, halting a run once ∼60x coverage has been achieved for 96 samples, and then washing and loading a new library with a further 96 samples offers further potential for cost savings. Sequencing 192 samples on a flow cell would reduce the cost to < £7 a genome using our approach.

By sequencing from single plaques we greatly reduced the time taken to obtain phage genomes, once a plaque is identified. Previously single plaque sequencing has been optimised for Illumina sequencing, by maximising the recovery of DNA from a single plaque and minimising the input into sequencing library preparation [38] . The previous approach does not scale well to the sequencing of hundreds or thousands of phage genomes, due to the filtration step and low input into Illumina sequencing libraries. The input for Illumina NexteraXT can be reduced to < 1 ng DNA when running an Illumina machine within an individual laboratory. However the capital costs of such machines are high and commercial suppliers do not routinely accept such low concentration of DNA, due to their quality control processes. Recently nanopore sequencing has been used to sequence single phage genomes from as little 0.4 ng DNA [39]. However, while feasible the approach is not practical for large-scale genome sequencing due to the limited number of reads that were obtained within a 72 hr run on a minION. While sufficient to sequence a single genome, the output from such low input does not allow rapid sequencing of hundreds of genomes [39].

To overcome the limited DNA obtained from a plaque, MDA was utilised. The use of MDA within our approach offers both advantages and disadvantages to genome sequencing. The primary advantage is the rapid increase in DNA that allows high-throughput approaches to be adopted. Secondary to this, it enables the sequencing of ssDNA phages, such as filamentous phages belonging to the family *Inoviridae*. MDA results in the dsDNA product from the ssDNA template, which can then be used to construct a sequencing library using a common, dsDNA-specific tagmentation approach. Finally, it also removes DNA modifications from bacteriophage genomes, which are known to be extensively hypermodified and recalcitrant to standard sequencing approaches [40–42]. The quantification of DNA after MDA allows the putative identification of phages which may have modified DNA that the EquiPhi enzyme cannot process, and be investigated further. The failure to produce amplified DNA, can be from lack of DNA input or modified DNA preventing amplification. The lack of output DNA may result from isolating phages with RNA genomes and thus have no DNA to amplify or phages with DNA modifications also preventing amplification.

The drawback of MDA is the removal of these biologically interesting DNA modifications, as they are replaced with standard nucleotides during amplification, so any modifications can no longer be detected or quantified. Additionally, amplification also removes the ability to precisely define the termini of phage genomes with approaches such as PhageTerm [43]. It should be noted the majority of phage genomes sequenced to date have utilised Illumina sequencing with NexteraXT [3] ,which also prevents recovery of genomic termini [43].

The use of MDA is known to introduce chimeras into reads [23]. The use of S1 nuclease did significantly reduce the number of chimeric reads, but the percentage of chimeric reads still remained high (48.5 and 49.9 in S1 nuclease treated samples). Thus, identification and splitting of these reads was necessary with SACRA [23]. It became apparent that the limiting step in obtaining a complete genome was the time for identification of chimeric reads with SACRA. SACRA functions by the identification of partially aligned reads compared to continuously aligned reads, and splitting of reads at a chimeric junction [23]. Although utilising SACRA provided an improvement in the number of assembled genomes, it proved to be a time-intensive step. To speed up the process of genome assembly, we took the approach of naively splitting reads into 300 bp fragments and treating them as short reads. This approach reliably increased the number of phage genomes, which is consistent with previous studies which showed that the majority of phage genomes can be assembled from short reads [35].

We incorporated a DNase I step to test the effectiveness of reducing bacterial DNA. With the *Pseudomonas* phage set there was no significant difference in the levels of host DNA when DNase I was used and was high across all samples. The presence of prophages which may be induced would increase the level of host contamination. To account for this we identified prophages with geNomad and manually removed them from the host genome and calculated the percentage of mapped reads to infer host DNA contamination. The predicted percentage of mapped reads still remained very high. There are likely numerous reasons for this. These include 1) not all prophages will be predicted and any remaining that are induced will be counted as host DNA using a mapping approach, 2) mapping allows sequence divergence, thus a closely related temperate phage that was isolated, will have reads that map to unidentified prophages in the host chromosome. 3) As phages are capable of both generalised and specialised transduction, which will package host DNA [44, 45]. We were unable to completely rule out that the different frequencies of transduction were not contributing to differences in the levels of host DNA. Despite the presence of host DNA within samples, it was still possible to assemble phages from samples, due to the relatively large amounts of data obtained. When DNAse I was used with *E. coli* MG1655, which does not contain an inducible prophage and the levels of host DNA are easily quantifiable, the percentage of DNA in samples was low ( < 3%) and did not hinder genome assembly. The levels of host DNA contamination in the cyanophage samples and range of hosts from the “Sheffield phages” was also very low compared to the *Pseudomonas* samples. We were unable to determine if DNase I was the reason for this. However, its addition does not prevent phage genome assembly. The heating of the sample to denature the DNase I, which also releases viral DNA from capsids at the same time does not seem to be problematic for low input amplifications, as seen by the successful genome sequencing of phages.

The development of a high-throughput, cost-effective approach, underpins future progress to make phage therapy a viable alternative or complement to antibiotics. Recent studies have demonstrated the predictive power of using phage genomes and host genomes to select phage that will infect [46, 47]. With the increasing size of datasets, the accuracy of such predictions will also increase. In selecting phages for therapy, it is desirable to know very early on in the selection of phages, if they contain genes which encode for antibiotic resistance genes or virulence factors [48], to prevent detailed characterisation of such phages. There have been several recent advances in the ability to rapidly isolate phages for therapeutic use [49, 50]. The approach described here now allows genome sequencing to now keep pace with that of isolation at greatly reduced cost.

## Conclusions

We have developed a cheap and rapid method which will facilitate the sequencing of hundreds of phages at greatly reduced cost. In demonstrating the approach, we showed it can be used to sequence phage genomes which vary in size from ∼5–279 kb, can be dsDNA or ssDNA, and works on plaques obtained from both Gram-positive and -negative bacterial hosts. Within the context of phage therapy it will enable the rapid development of robust phage biobanks, built upon genomic data. The reduced cost per genome and low capital costs, allows phagebanks to be developed in situ in low and middle income countries, where the need for phage biobanks is often greatest. The sequencing of single plaques on *E. coli*, also identified only 4 of 96 plaques contained multiple phages, demonstrating it is not necessary to utilise three rounds of plaque purification to obtain purified phages for further characterisation. Within a broader context it will allow for a widespread expansion in both the number and diversity of phage genomes, which are greatly under-represented compared to their bacterial hosts.

## Data availability

Genomic data was submitted to the project accessions PRJEB94355 and PRJEB89903. Code is available from https://github.com/amillard/plaque2seq.

## Funding Statements

This work was supported by LifeArc and Cystic Fibrosis Trust under grant no. THUB03 for the Trailfinder-CF Innovation Hub, as part of the Translational Innovation Hub Network for CF Lung Health & Infection. LifeArc is registered as a charity in England and Wales (1015243) and in Scotland (SC037861). Cystic Fibrosis Trust is registered as a charity in England and Wales (1079049) and in Scotland (SC040196). GNM Jr. and DJS were funded by the European Research Council (Grant agreement ID 883551). RJP was funded by NERC (Grant ref. NE/Z00019X/1). ES is funded by the NIHR Sheffield Biomedical Research Centre (BRC) (NIHR203321). ADM was supported by MRC (MR/T030062/1). AP, TJ and JCDL were supported by BBSRC Midlands Integrative Biosciences Training Partnership (MIBTP) DTP

## Author Contribution

A.M. and S.M. conceived the study. A.M. , S. M and A.K. developed the code. A.M., S.M., R.G., and A.K. analysed the data. A.M., S.M., and R.G. drafted the manuscript. E.G., M.R.J.C., R.J.P., D.J.S., G.S., and W.v.S. supervised the work. Data were generated by R.G., A.K., K.P., K.W., S.H., N.U., A.A., C.E., S.R., J.C.D.L., F.P., T.J., A.P., M.C., H.F.Y.A.-D., G.A.S., N.S.R., H.S., and G.N.M.

---

**Figure S1.**
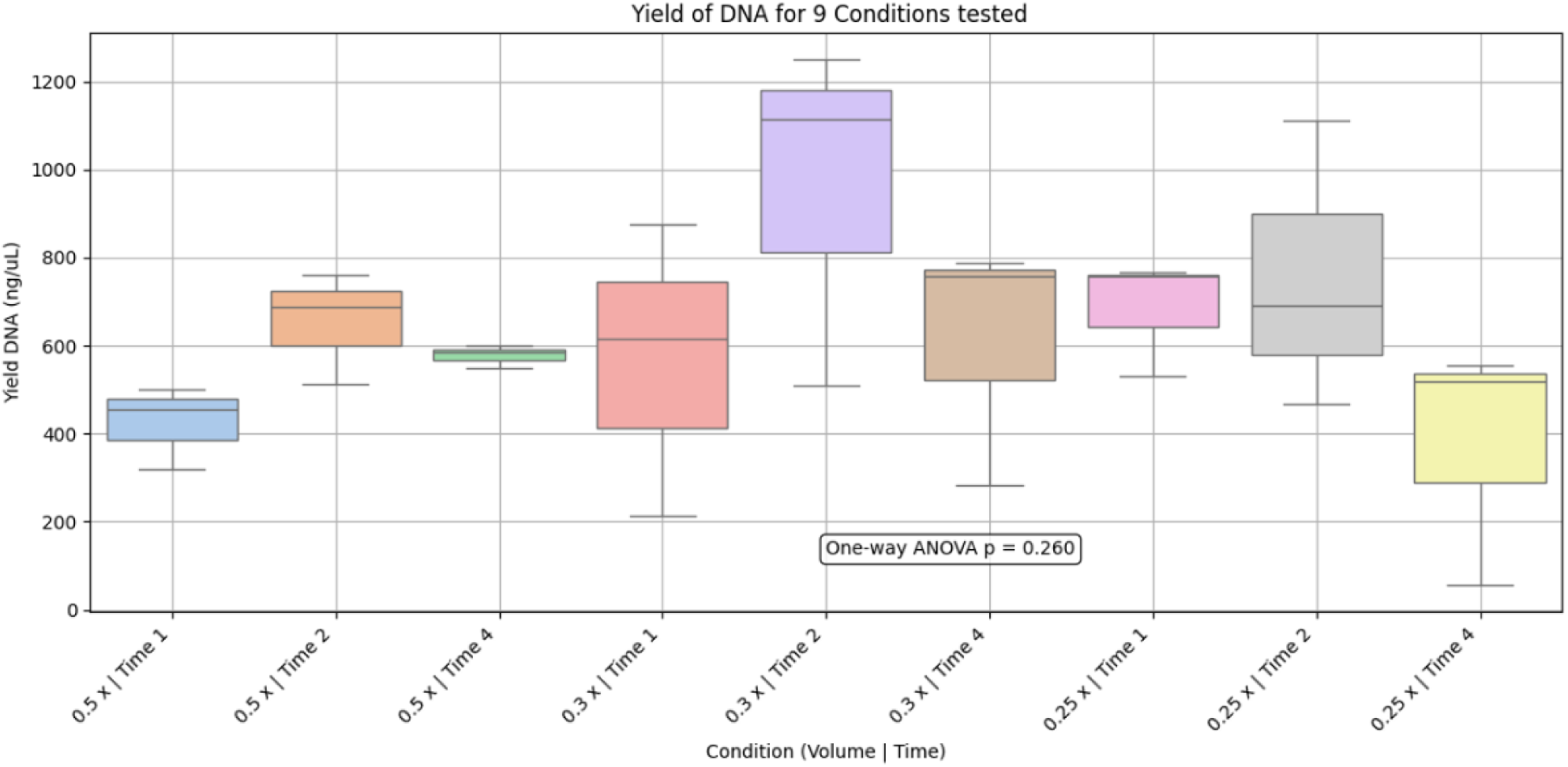
Multiple displacement amplification of DNA from plaques. Twenty seven plaques were resuspended in SM buffer and MDA applied. The following conditions were applied, amplification for 1, 2 and 4 hours and amplification using 0.5x , 0.3x and 0.25x the standard 20 ul volumes. To give a total of 9 conditions, with 3 replicates per condition. No significant difference was found for DNA yield, between conditions ( one way ANOVA).

**Figure S2.**
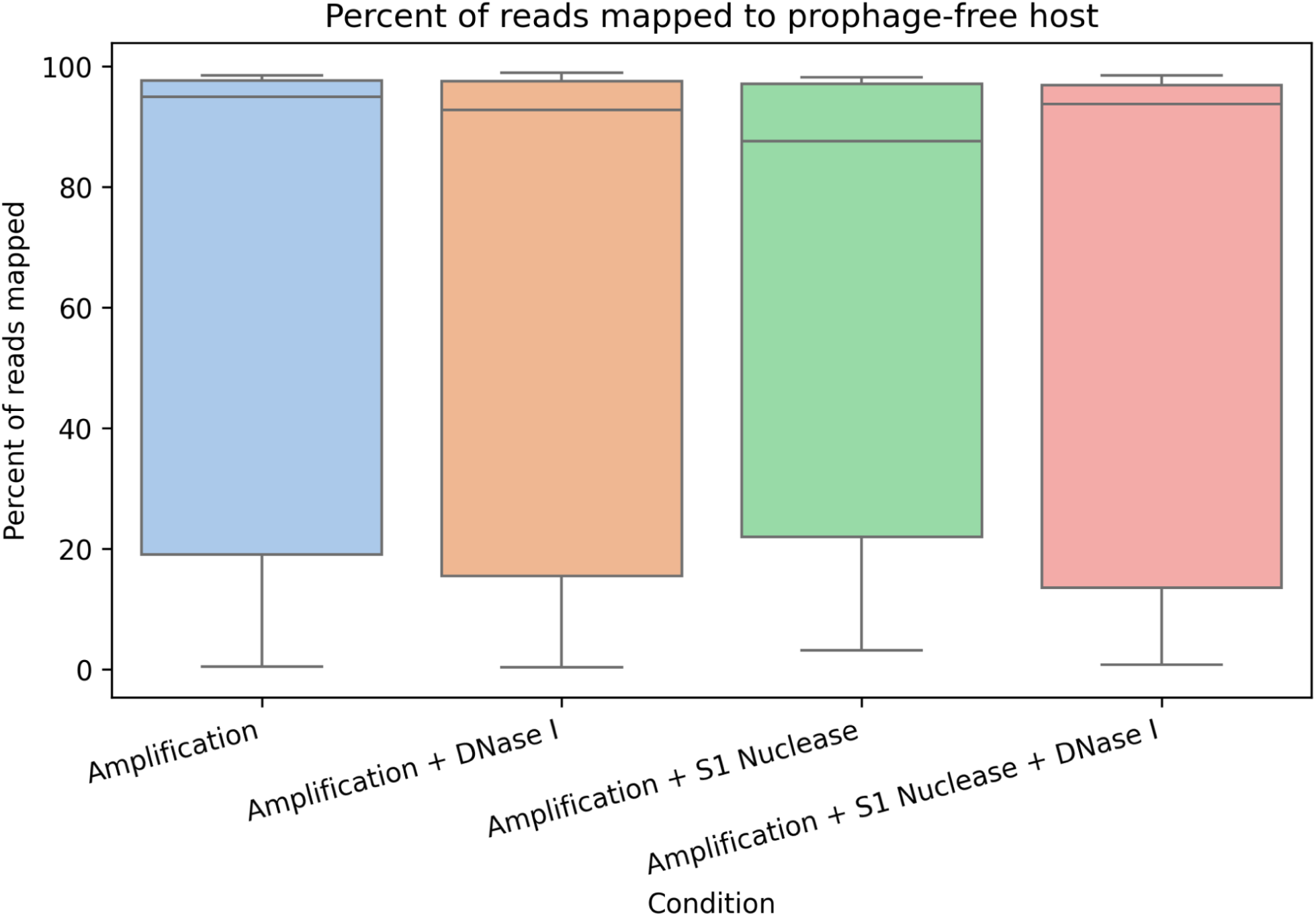
Detection of host DNA in four treatment groups. The percentage of host reads was determined by mapping of reads against the *Pseudomonas* host, which had predicted prophages removed. Data plotted has been filtered to only include samples where sequence was obtained in all four treatments, < 100% of reads mapped to the host (to excluded samples with no phage) and >10000 reads (removal of samples with low sequencing output). Treatment 1:MDA only S1 Nuclease; Treatment 2: MDA and DNase I; Treatment 3: MDA and S1 Nuclease; Treatment 4: MDA, S1 Nuclease and DNase I.

**Figure S3.**
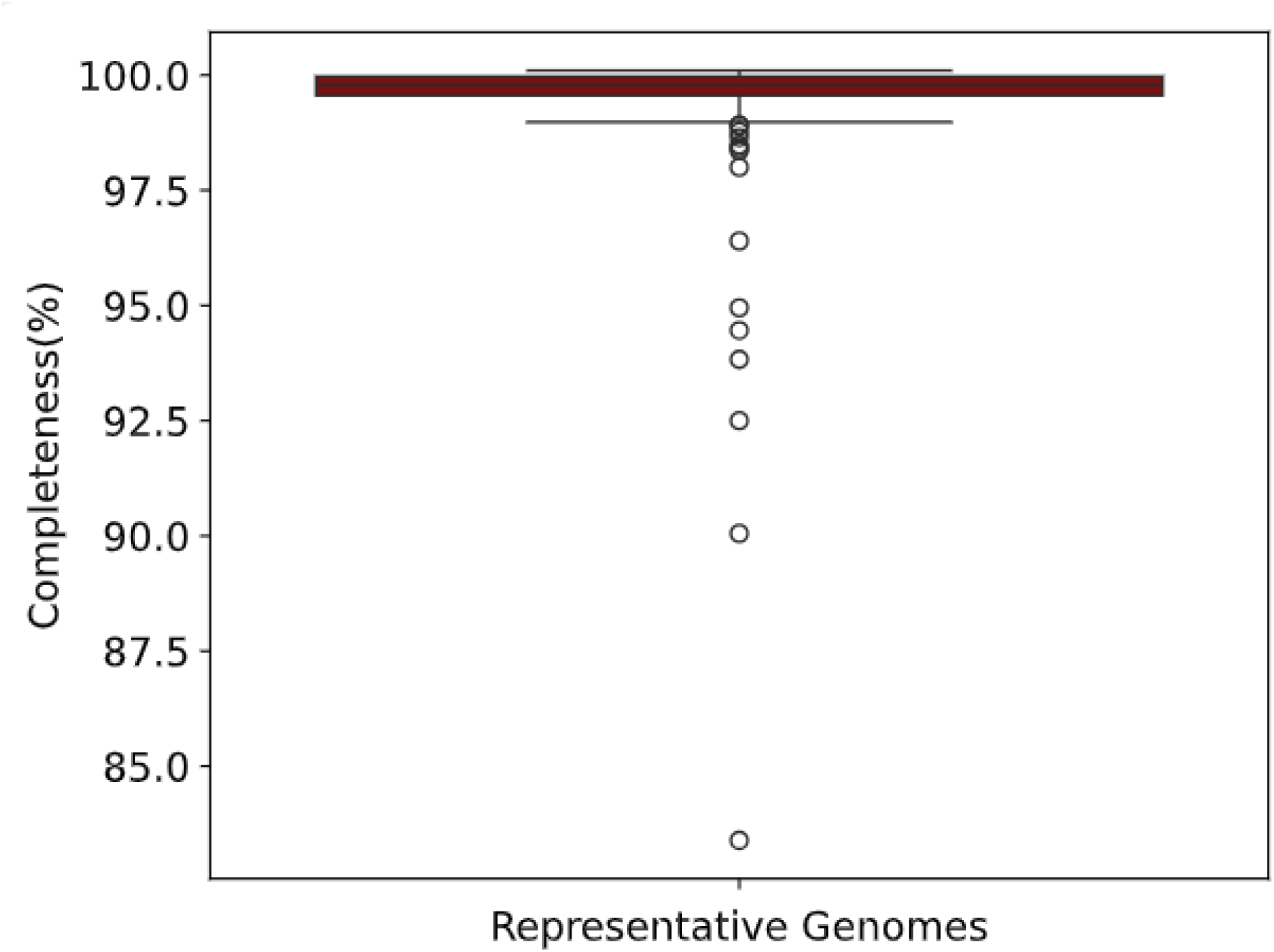
Assembly of *in silico* long reads converted to short reads. In silico long reads were generated for a set of 197 diverse phage genomes using PBSIM [19], converted into short 300 bp reads and assembled with SPades. Genome completeness was determined by comparison to the original genome sequence length.

**Figure S4.**
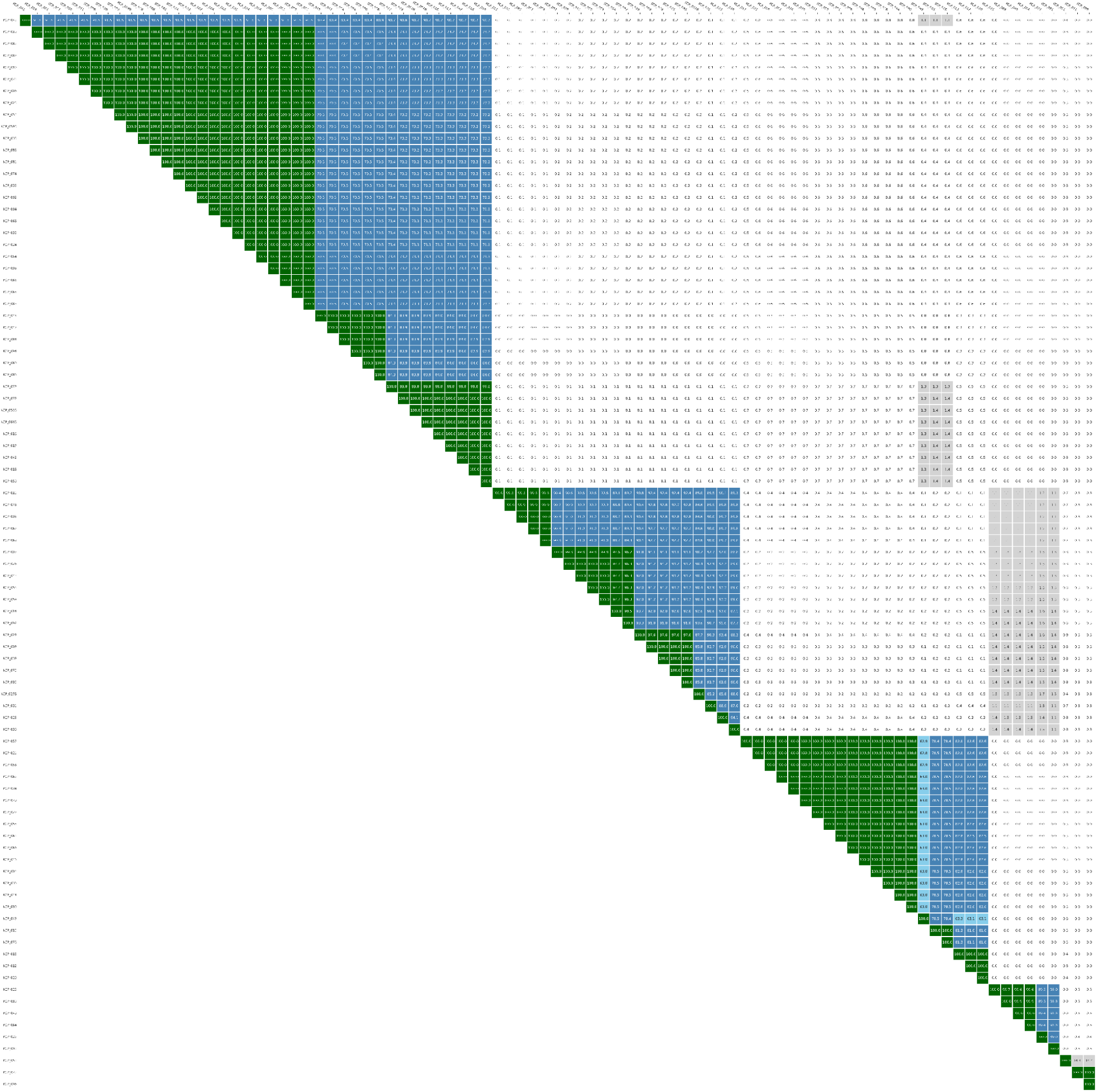
Comparative Genomics of Coliphages. All 95 coliphages were compared using *taxMyPhage* using the similarity option.

**Figure S5.**
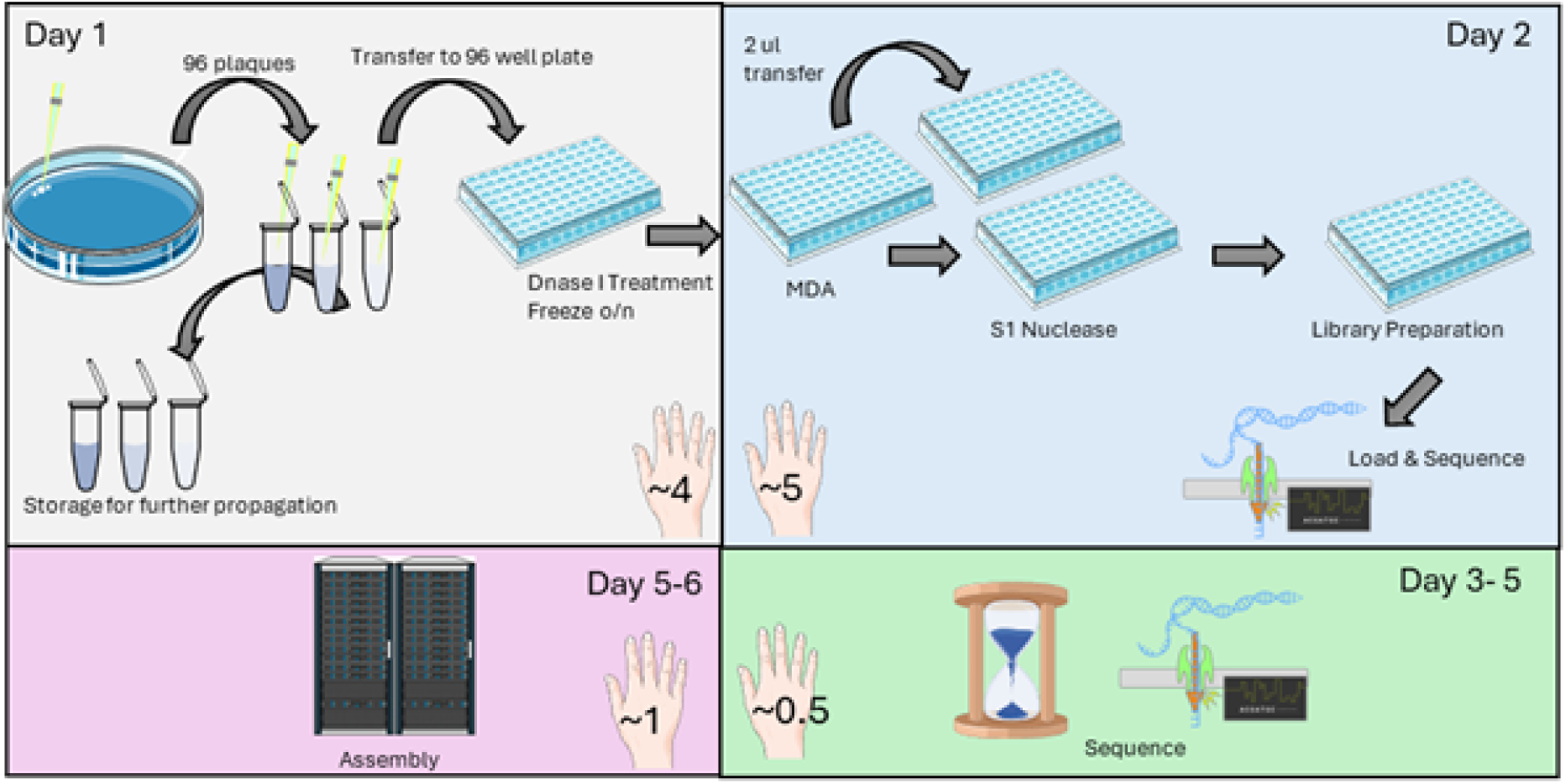
Plaque-2-sequence workflow overview. Overview of optimised pipeline for genome sequence. Day 1: plaques are picked into 100 μl of SM buffer and left for 1 hr. 10 μl is transferred to 96 well PCR plate and DNAse I added, to degrade free host DNA, prior to heat denaturation at 95 °C. Samples are stored frozen overnight. Day 2: samples amplified with MDA, prior to S1 nuclease treatment to reduce chimeric DNA. Rapid Barcoding kit V96 14 is used for library preparation, followed by loading onto a minION flow cell. Days 2–5: sequencing occurs. Day 5–6:assembly of genomes.

## Notes

### Competing Interest Statement

The authors have declared no competing interest.

## Reference list

1. E. White H, V. Orlova E (2020) Bacteriophages: Their structural organisation and function. Bacteriophages - Perspectives and Future

2. Dion MB, Oechslin F, Moineau S (2020) Phage diversity, genomics and phylogeny. Nat Rev Microbiol 18:125–138

3. Cook R, Brown N, Redgwell T, Rihtman B, Barnes M, Clokie M, Stekel DJ, Hobman J, Jones MA, Millard A (2021) INfrastructure for a PHAge REference Database: Identification of large-scale biases in the current collection of cultured phage genomes. PHAGE. 10.1089/phage.2021.0007

4. Turner D, Shkoporov AN, Lood C, et al (2023) Abolishment of morphology-based taxa and change to binomial species names: 2022 taxonomy update of the ICTV bacterial viruses subcommittee. Arch Virol 168:74

5. Oksanen HM, Ictv Report Consortium (2017) ICTV virus taxonomy profile: Corticoviridae. J Gen Virol 98:888–889

6. Knezevic P, Adriaenssens EM, Ictv Report Consortium (2021) ICTV virus Taxonomy profile: Inoviridae. J Gen Virol. 10.1099/jgv.0.001614

7. Wu Y, Wu Z, Guo L, Shao J, Xiao H, Yang M, Deng C, Zhang Y, Zhang Z, Zhao Y (2024) Diversity and distribution of a prevalent Microviridae group across the global oceans. Commun Biol 7:1377

8. Callanan J, Stockdale SR, Adriaenssens EM, Kuhn JH, Rumnieks J, Pallen MJ, Shkoporov AN, Draper LA, Ross RP, Hill C (2021) Leviviricetes: expanding and restructuring the taxonomy of bacteria-infecting single-stranded RNA viruses. Microb Genom. 10.1099/mgen.0.000686

9. Adriaenssens EM, Cowan DA (2014) Using signature genes as tools to assess environmental viral ecology and diversity. Appl Environ Microbiol 80:4470–4480

10. Cook R, Hooton S, Trivedi U, King L, Dodd CER, Hobman JL, Stekel DJ, Jones MA, Millard AD (2021) Hybrid assembly of an agricultural slurry virome reveals a diverse and stable community with the potential to alter the metabolism and virulence of veterinary pathogens. Microbiome 9:65

11. Zaragoza-Solas A, Haro-Moreno JM, Rodriguez-Valera F, López-Pérez M (2022) Long-read metagenomics improves the recovery of viral diversity from complex natural marine samples. mSystems e0019222

12. Elek CKA, Brown TL, Le Viet T, et al (2023) A hybrid and poly-polish workflow for the complete and accurate assembly of phage genomes: a case study of ten przondoviruses. Microb Genom. 10.1099/mgen.0.001065

13. Pirnay J-P, Blasdel BG, Bretaudeau L, et al (2015) Quality and safety requirements for sustainable phage therapy products. Pharm Res 32:2173–2179

14. Nagel T, Musila L, Muthoni M, Nikolich M, Nakavuma JL, Clokie MR (2022) Phage banks as potential tools to rapidly and cost-effectively manage antimicrobial resistance in the developing world. Curr Opin Virol 53:101208

15. Yukgehnaish K, Rajandas H, Parimannan S, Manickam R, Marimuthu K, Petersen B, Clokie MRJ, Millard A, Sicheritz-Pontén T (2022) PhageLeads: Rapid Assessment of Phage Therapeutic Suitability Using an Ensemble Machine Learning Approach. Viruses. 10.3390/v14020342

16. Bouras G, Nepal R, Houtak G, Psaltis AJ, Wormald P-J, Vreugde S (2023) Pharokka: a fast scalable bacteriophage annotation tool. Bioinformatics. 10.1093/bioinformatics/btac776

17. Kropinski AM, Mazzocco A, Waddell TE, Lingohr E, Johnson RP (2009) Enumeration of bacteriophages by double agar overlay plaque assay. Methods Mol Biol 501:69–76

18. dorado: Oxford Nanopore’s Basecaller. Github

19. Ono Y, Hamada M, Asai K (2022) PBSIM3: a simulator for all types of PacBio and ONT long reads. NAR Genom Bioinform 4:lqac092

20. Bushnell B (2014) BBMap: A fast, accurate, splice-aware aligner. Lawrence Berkeley National Lab. (LBNL), Berkeley, CA (United States)

21. Bankevich A, Nurk S, Antipov D, et al (2012) SPAdes: A New Genome Assembly Algorithm and Its Applications to Single-Cell Sequencing. J Comput Biol 19:455–477

22. Kurtz S Phillippy A DALSMSMACSSL (2004) Versatile and open software for comparing large genomes. Genome Biol.

23. Kiguchi Y, Nishijima S, Kumar N, Hattori M, Suda W (2021) Long-read metagenomics of multiple displacement amplified DNA of low-biomass human gut phageomes by SACRA pre-processing chimeric reads. DNA Res. 10.1093/dnares/dsab019

24. Nayfach S, Camargo AP, Schulz F, Eloe-Fadrosh E, Roux S, Kyrpides NC (2021) CheckV assesses the quality and completeness of metagenome-assembled viral genomes. Nat Biotechnol 39:578–585

25. Millard A, Denise R, Lestido M, Thomas MT, Webster D, Turner D, Sicheritz-Pontén T (2025) TaxMyPhage: Automated taxonomy of dsDNA phage genomes at the genus and species level. Phage (New Rochelle). 10.1089/phage.2024.0050

26. Shen A, Millard A (2021) Phage genome annotation: Where to begin and end. PHAGE 2:183–193

27. pysam. In: https://github.com/pysam-developers/pysam.

28. (2023) Fast and accurate identification of plasmids and viruses in sequencing data using geNomad. Nat Biotechnol. 10.1038/s41587-023-01982-7

29. Terzian P, Olo Ndela E, Galiez C, Lossouarn J, Pérez Bucio RE, Mom R, Toussaint A, Petit M-A, Enault F (2021) PHROG: families of prokaryotic virus proteins clustered using remote homology. NAR Genom Bioinform 3:lqab067

30. Wilson WH, Carr NG, Mann NH (1996) The effect of phosphate status on the kinetics of cyanophage infection in the oceanis cyanobacterium Synechococcus sp. WH7803. J Phycol 32:506–516

31. Seemann T (2014) Prokka: Rapid prokaryotic genome annotation. Bioinformatics 30:2068–2069

32. Vaser R, Šikić M (2021) Time- and memory-efficient genome assembly with Raven. Nat Comput Sci 1:332–336

33. Li H (2018) Minimap2: pairwise alignment for nucleotide sequences. Bioinformatics 34:3094–3100

34. Mehta P, Casjens S, Krishnaswamy S (2004) Analysis of the lambdoid prophage element e14 in the E. coli K-12 genome. BMC Microbiol 4:4

35. Rihtman B, Meaden S, Clokie MRJ, Koskella B, Millard AD (2016) Assessing Illumina technology for the high-throughput sequencing of bacteriophage genomes. PeerJ 4:e2055

36. Kurtz S, Phillippy A, Delcher AL, Smoot M, Shumway M, Antonescu C, Salzberg SL (2004) Versatile and open software for comparing large genomes. Genome Biol 5:R12

37. Wang Y, Zhao Y, Bollas A, Wang Y, Au KF (2021) Nanopore sequencing technology, bioinformatics and applications. Nat Biotechnol 39:1348–1365

38. Kot W, Vogensen FK, Sørensen SJ, Hansen LH (2014) DPS - A rapid method for genome sequencing of DNA-containing bacteriophages directly from a single plaque. J Virol Methods 196:152–156

39. Sbaghdi T, Jagorel F, Monot M, Garneau JR (2025) Short-read and long-read PCR-free sequencing of bacteriophages using ultra-low starting DNA input. J Biomol Tech. 10.7171/3fc1f5fe.c0001573

40. Lee Y-J, Dai N, Walsh SE, Müller S, Fraser ME, Kauffman KM, Guan C, Corrêa IR, Weigele PR (2018) Identification and biosynthesis of thymidine hypermodifications in the genomic DNA of widespread bacterial viruses. Proceedings of the National Academy of Sciences 201714812

41. Puxty RJ, Millard AD (2023) Functional ecology of bacteriophages in the environment. Curr Opin Microbiol 71:102245

42. Rihtman B, Puxty RJ, Hapeshi A, et al (2021) A new family of globally distributed lytic roseophages with unusual deoxythymidine to deoxyuridine substitution. Curr Biol 31:3199–3206.e4

43. Garneau J, Depardieu F, Fortier L-C, Bikard D, Monot M (2017) PhageTerm : a Fast and User-friendly Software to Determine Bacteriophage Termini and Packaging Mode using randomly fragmented NGS data. Sci Rep 7:7

44. Chen J, Quiles-Puchalt N, Chiang YN, Bacigalupe R, Fillol-Salom A, Chee MSJ, Ross Fitzgerald J, Penadés JR (2018) Genome hypermobility by lateral transduction. Science 362:207–212

45. Matilla M a., Fang X, Salmond GP (2014) Viunalikeviruses are environmentally common agents of horizontal gene transfer in pathogens and biocontrol bacteria. ISME J 1–5

46. Gaborieau B, Vaysset H, Tesson F, et al (2024) Prediction of strain level phage-host interactions across the Escherichia genus using only genomic information. Nat Microbiol 9:2847–2861

47. Keith M, Park de la Torriente A, Chalka A, Vallejo-Trujillo A, McAteer SP, Paterson GK, Low AS, Gally DL (2024) Predictive phage therapy for Escherichia coli urinary tract infections: Cocktail selection for therapy based on machine learning models. Proc Natl Acad Sci U S A 121:e2313574121

48. Cook BWM, Hynes AP (2025) Re-evaluating what makes a phage unsuitable for therapy. NPJ Antimicrob Resist 3:45

49. Nair G, Chavez-Carbajal A, Tullio RD, et al (2024) Micro-plaque assays: A high-throughput method to detect, isolate, and characterize bacteriophages. bioRxiv. 10.1101/2024.06.20.599855

50. Olsen NS, Hendriksen NB, Hansen LH, Kot W (2020) A New High-Throughput Screening Method for Phages: Enabling Crude Isolation and Fast Identification of Diverse Phages with Therapeutic Potential. Phage (New Rochelle) 1:137–148

